# Disturbance-induced changes in size-structure promote coral biodiversity

**DOI:** 10.1101/2020.05.21.094797

**Authors:** Mariana Álvarez-Noriega, Joshua S. Madin, Andrew H. Baird, Maria Dornelas, Sean R Connolly

**Author notes:** Corresponding author: Mariana Álvarez-Noriega, Centre for Geometric Biology, School of Biological Sciences Monash University, Melbourne, VIC, Australia 3800.

## Abstract

Reef-building coral assemblages are typically species-rich, yet the processes maintaining coral biodiversity remain poorly understood. Disturbance has long been believed to promote coral species coexistence by reducing the strength of competition. However, such disturbance-induced effects have since been shown to be insufficient on their own to prevent competitive exclusion. Nevertheless, Modern Coexistence Theory has revealed other mechanisms by which disturbance and, more generally, environmental variation can favour coexistence. Here, we formulate, calibrate, and analyze a size-structured, stochastic coral competition model using field data from two common colony morphologies. These two coral morphologies, tabular and digitate, differ in their size-dependent vulnerability to dislodgement caused by wave action. We confirm that fluctuations in wave action can promote coral species coexistence. However, using a recently proposed partitioning framework, we show that, contrast to previous expectations, temporal variability in strength of competition did not promote coexistence. Instead, coexistence was enabled by differential fluctuations in size-dependent mortality among competitors. Frequent and intense disturbances resulted in monocultures of digitate corals, which are more robust to wave action than tabular corals. In contrast, infrequent or weak disturbances resulted in monocultures of tabular corals. Coexistence was only possible under intermediate levels of disturbance frequency and intensity. Given the sensitivity of coexistence to disturbance frequency and intensity, anthropogenic changes in disturbance regimes are likely to affect biodiversity in coral assemblages in ways that are not predictable from single population models.

## Introduction

Reef-building coral assemblages are an example of the ‘paradox of the plankton’ (Hutchinson 1961): they can be species-rich, even though species compete for a small number of limiting resources, mainly space, light, and nutrients in the water column. However, the processes maintaining such high biodiversity remain enigmatic (Tanner *et al.* 1994; Bellwood & Hughes 2001). A classical explanation for reef coral coexistence, the intermediate disturbance hypothesis (IDH), involves the periodic reduction, by disturbances, in densities of dominant competitors, which frees up space for colonisation by weaker competitors (Connell 1978; Aronson & Precht 1995). However, the IDH’s theoretical validity has been challenged because the weakening of competition in the presence of disturbance is not sufficient to promote coexistence (Chesson & Huntly 1997; Fox 2013). Long-term coexistence requires that competition operates in such a way that individuals experience progressively less competition, on average, as they become rare, so that they tend to recover from excursions to low density. While disturbance cannot promote coexistence solely by weakening competition, it produces environmental fluctuations that under some circumstances can promote coexistence (Roxburgh *et al.* 2004).

Environmental fluctuations can favour coexistence when population growth rate responds nonlinearly or sub-additively to these environmental fluctuations or competition. A sub-additive response to the environment and competition means that population growth rate is reduced less by competition when a species is experiencing an unfavourable environment than when it is experiencing a favourable one. The higher cost of competition during favourable times acts as an upper bound to population growth, while the lower cost during unfavourable times acts as a lower bound. The coexistence mechanism acting via competitive and environmental sub-additivity is called the *storage effect* (Chesson & Warner 1981; Chesson 2000), which is known to operate in many ecological assemblages (e.g., (Cáceres 1997; Adler *et al.* 2006; Angert *et al.* 2009)). However, covariation between other factors or demographic rates affecting population growth can also impact coexistence (Ellner *et al.* 2019).

When population growth rate responds nonlinearly to competition, the average population growth can either be boosted or depressed by fluctuations in competition relative to a constant environment at the mean competition (Armstrong & McGehee 1980) (i.e., *relative nonlinearity of competition;* (Chesson 2000)). Coexistence is possible when the inferior competitor under average conditions gets a larger benefit from fluctuations in competition than does the superior competitor, and each population, when abundant, creates the conditions that favour its competitor. Coexistence via nonlinear averaging has not received as much attention as the storage effect, despite being more important than non-additivity under some conditions (e.g., (Miller *et al.* 2011; Letten *et al.* 2018)). Moreover, until very recently, coexistence via nonlinear averaging was thought to act exclusively via nonlinearities in the population growth rate’s response to competition, however nonlinear responses to other factors, such as the environment, can also promote coexistence (Ellner *et al.* 2019).

In coral assemblages, hydrodynamic disturbances strongly affect assemblage structure (Connell *et al.* 2004), mainly by imposing mortality pulses that affect top-heavy colonies more than bottom-heavy ones (Madin & Connolly 2006a). Consequently, susceptibility to wave action is morphology- and size-dependent (Massel & Done 1993; Madin & Connolly 2006a; Madin *et al.* 2014). Since top-heavy colonies, such as those of species with a tabular morphology, tend to grow faster than bottom heavy colonies (Dornelas *et al.* 2017) and can monopolize space on the reef crest in the absence of hydrodynamic disturbances (Baird & Hughes 2000), periodic hydrodynamic disturbances have long been thought to prevent the exclusion of bottom-heavy competitors (Connell 1978). Moreover, because top-heavy colonies increase in susceptibility to mechanical dislodgement with increasing colony size, storms affect the largest, most fecund sizes disproportionally, further affecting superior competitors. Therefore, the effect of hydrodynamic disturbances on coexistence acts in two ways: clearing space for larvae to settle and altering, not just the relative number of colonies of each competitor, but also the populations’ size structures. While the presence of periodic disturbance is not sufficient to promote coexistence (Chesson & Huntly 1997; Fox 2013), if a competitors’ population growth rates respond nonlinearly or non-additively to factors (e.g., competition) or demographic rates (e.g., survival) that fluctuate with disturbance, disturbance could promote coexistence between coral morphologies.

Here, we calibrated a competition model using field demographic data for two common coral morphologies (Madin *et al.* 2014; Álvarez-Noriega *et al.* 2016; Dornelas *et al.* 2017) and simulated hydrodynamic disturbances using a local 37-year wind record. Using model simulations, we first investigated whether or not coexistence was possible in the presence of hydrodynamic disturbance. Then, having found that it was, we identified the fluctuation-dependent mechanisms responsible for coexistence by decomposing competitors’ population growth rates into contribution from each of the different fluctuating components: competition, size structure, and the interaction between the two. We chose tabular *Acropora* as the model morphology for corals that are fast growing, good competitors that are highly susceptible to disturbance, and digitate *Acropora* as the model morphology for slower growing, more mechanically robust corals. These two morphologies are very common in the wave-exposed habitats of highly diverse Indo-Pacific reefs (Done 1982; Dornelas & Connolly 2008).

## Results and Discussion

In our model, coexistence of tabular and digitate corals was only possible in the presence of hydrodynamic disturbance (Fig. 1). In a variable environment, both competitors had a positive population growth rate when they were rare and the other competitor was a resident (Fig. 1-a & b). This implies that if the population of either competitor reaches very low densities, it will be able to recover and avoid extinction in the presence of the other competitor, which is abundant (or at least at its long-term abundance in monoculture). However, in a constant environment, one of the competitors was unable to recover from low densities (Fig. 1-c & d). Indeed, with our model, we find no region of parameter space where coexistence could occur in the absence of fluctuations (see Methods). To investigate the mechanism by which hydrodynamic disturbance was driving coexistence, we isolate the effects of the different sources of fluctuations in our model on invader growth rate. We quantified the approximate contribution of the two main fluctuating factors affected by wave action: competition (i.e., proportion of free space) and size-dependent mortality due to disturbance, as well as the additional contribution of both factors fluctuating together (i.e., their interaction). To do this, we used a recent quantitative framework (Ellner *et al.* 2019) that relies on simulations rather than mathematical approximations, thus allowing for higher model complexity than previous frameworks (e.g.,(Chesson 1994)).

**Figure 1.**
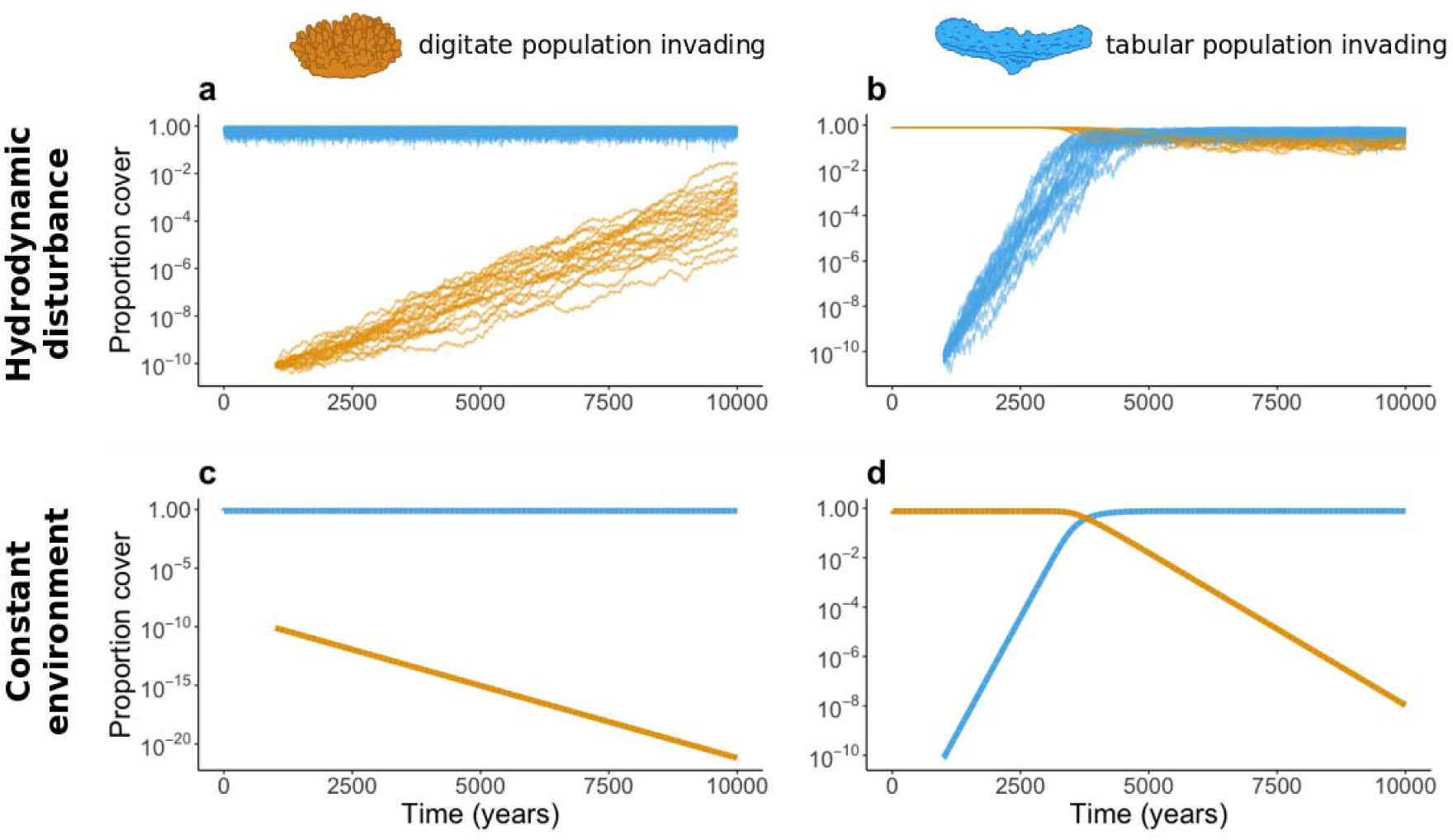
Proportion of space occupied through time following an invasion. The trajectories of the digitate and tabular populations are in orange and blue, respectively. Panel a- Digitate population invading a tabular resident in the presence of environmental fluctuations. Panel b- Tabular population invading a digitate resident in the presence of environmental fluctuations. Panel c- Digitate population invading a tabular resident at the mean size-dependent survival rates (constant environment). Panel d- Tabular population invading a digitate resident at the mean size-dependent survival rates (constant environment). Note that the scale is different in Panel c. In panels a & b, each line represents one simulation (out of 20), in panels c & d, all simulations follow the same trajectory. These simulations were run for 10,000 years (with invasion after 2000 years) to show the full dynamics of invasion, but were run only for 2300 years (300 years following invasion) for all other analyses.

Contrary to the prevailing understanding of how disturbance promotes coexistence, fluctuations in free space actually had a coexistence-inhibiting effect (i.e., a negative effect on the invader growth rate; Fig. 2-a, Fig. S6), relative to the constant free space model where average free space from the stochastic simulation was imposed every year. Rather, invasion and establishment of the digitate population depended on the beneficial effect of fluctuations in size structure caused by differences in size-dependent susceptibility to disturbance (which was the only term >0 for the digitate population; Fig. 2-a). The positive contribution of fluctuations in size structure counteracted the negative effects of fluctuations in free space and size structure varying jointly on the digitate population’s invasion growth rate.

**Figure 2.**
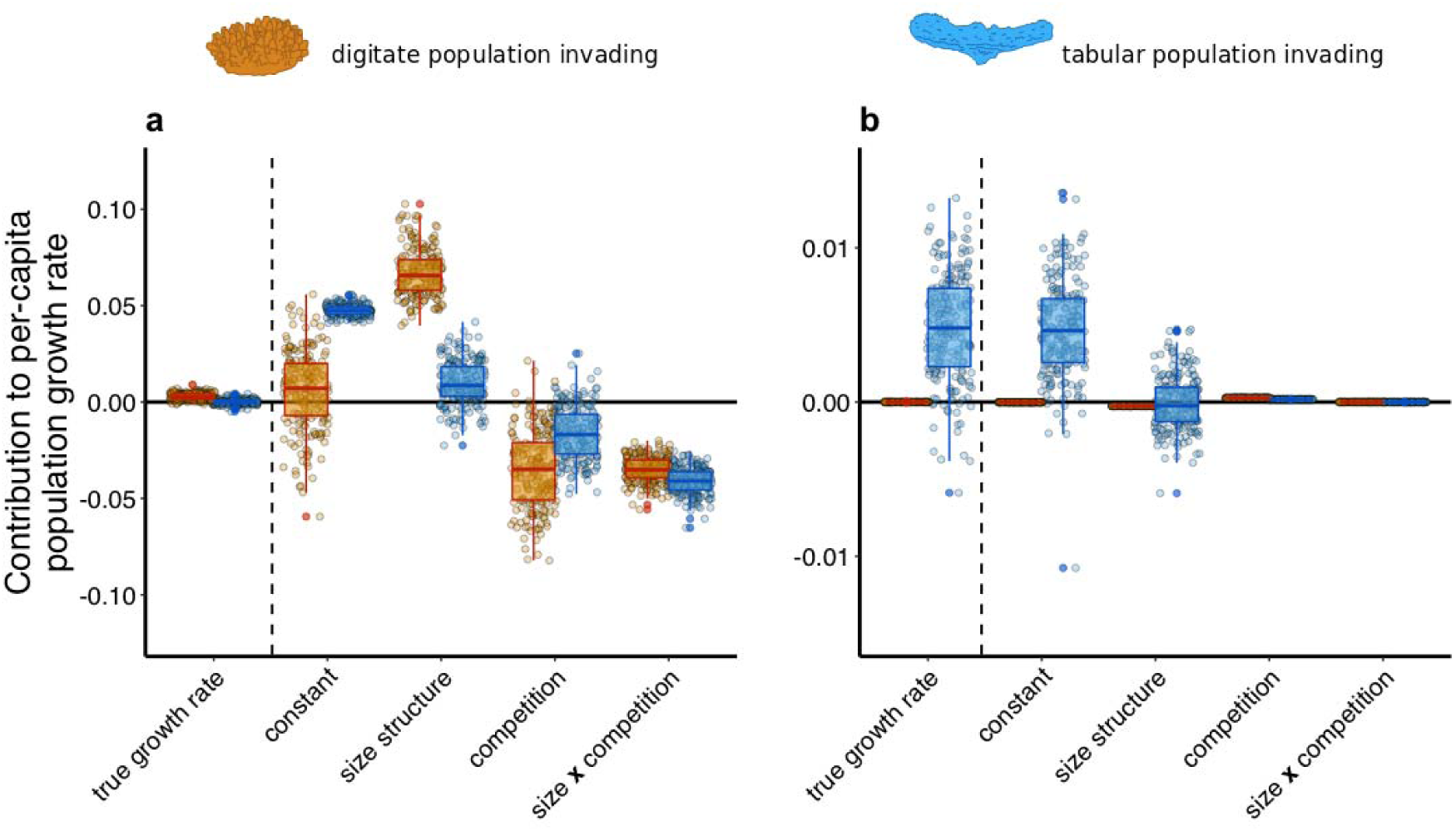
Partitioning the contributions to per-capita population growth (r). Contribution of the different sources of variation to mean population growth rate (‘true population growth’) following Ellner et al. 2019 (Ellner *et al.* 2019) when a digitate population invades a tabular resident (Panel a), and when a tabular population invades a digitate resident (Panel b). *Constant* refers to the population growth rate at mean values of free space and competitors’ size structures. *Size structure* represents the contribution of fluctuations in size structure to population growth rate when competition is constant (at its mean value). Similarl*y, competition* represents the contribution of fluctuations in the amount free space when size structures are fixed at their mean values for each competition. *Size structure* and *competition* are the main effects. *Size* **x** *competition* represents the contribution of the simultaneous fluctuation of competition and size structure that is independent of their main effects. The true population growth rate is the sum of *constant, size structure, competition*, and *size* **x** *competition*, and it is equal to the mean population growth rate when size structure and competition fluctuate (). See methods for more details. The digitate and tabular populations are represented by orange and blue colours, respectively. Each point represents the one simulation (200 simulations in total), and the box plots show their distribution.

Conversely, the invasion growth rate of the tabular population was positive solely because it had a positive per-capita growth rate under average conditions (Fig. 2-b). Since the digitate population was not vulnerable to hydrodynamic dislodgment, fluctuations were negligible when this population was dominant, and therefore did not affect population growth.

To understand how environmental fluctuations induced a stabilizing coexistence effect, it is important to recognize that each species was favoured at different levels of environmental variation. Hydrodynamic disturbance increased the digitate population’s growth rate, while the tabular population performed best when environmental variation was low (i.e., under average conditions). When abundant, each population created the conditions that favoured its competitor, allowing the latter to increase in abundance. When the digitate population was rare, the high abundance of tabular corals induced large fluctuations in competition – specifically, via the dislodgment of large tabular corals, and an associated reduction in reproductive output of the tabular population – which favoured the digitate population. Conversely, when the tabular population was rare and the mechanically-stable digitate population was abundant, resource fluctuations were minimal. This favoured tabular corals, whose per-capita population growth rate was higher at the mean resource level, as well as in the absence of disturbance. These dynamics are consistent with empirical observations. Tabular corals grow faster than digitate corals (Dornelas *et al.* 2017) and reach larger colony sizes that are very fecund (Álvarez-Noriega *et al.* 2016). In periods of low hydrodynamic disturbance, tabular species can dominate the reef crest (Baird & Hughes 2000). However, hydrodynamic disturbances affect the large, very fecund, tabular colonies most strongly (Madin & Connolly 2006a) and, consequently, the relative abundances of digitate corals tend to increase after disturbances that dislodge tabular corals (Muko *et al.* 2013). Our findings are also consistent with changes in the relative abundance between species following disturbance in the Caribbean (Aronson & Precht 1995) and the recovery of species richness following strong hydrodynamic disturbance on the GBR (Connell *et al.* 2004).

According to the IDH (Connell 1978), coexistence should be more likely at intermediate levels of disturbance because high disturbance eliminates susceptible species, while low disturbance allows dominant competitors to exclude inferior ones. Sensitivity analysis indicates that fluctuation-dependent coexistence creates such a relationship in our model: if disturbance is weak or very infrequent, the tabular population outcompetes the digitate population; if disturbance is very frequent, the digitate population outcompetes the tabular population (Fig. 3-a). Coexistence was possible for a wide range of disturbance frequencies and intensities but, as expected, the more intense the disturbance, the less frequent it could be and vice-versa. Once disturbance was sufficiently intense to dislodge most of the digitate colonies (i.e. only the largest sizes would survive; Fig. 3-b), the digitate population was again unable to invade a tabular resident, despite most tabular colonies being dislodged too (Fig. 3-a & b). This occurred because the rapid growth of surviving small tabular colonies favoured their population recovery over that of digitate populations, for extreme disturbance regimes that dislodged even digitate colonies.

**Figure 3.**
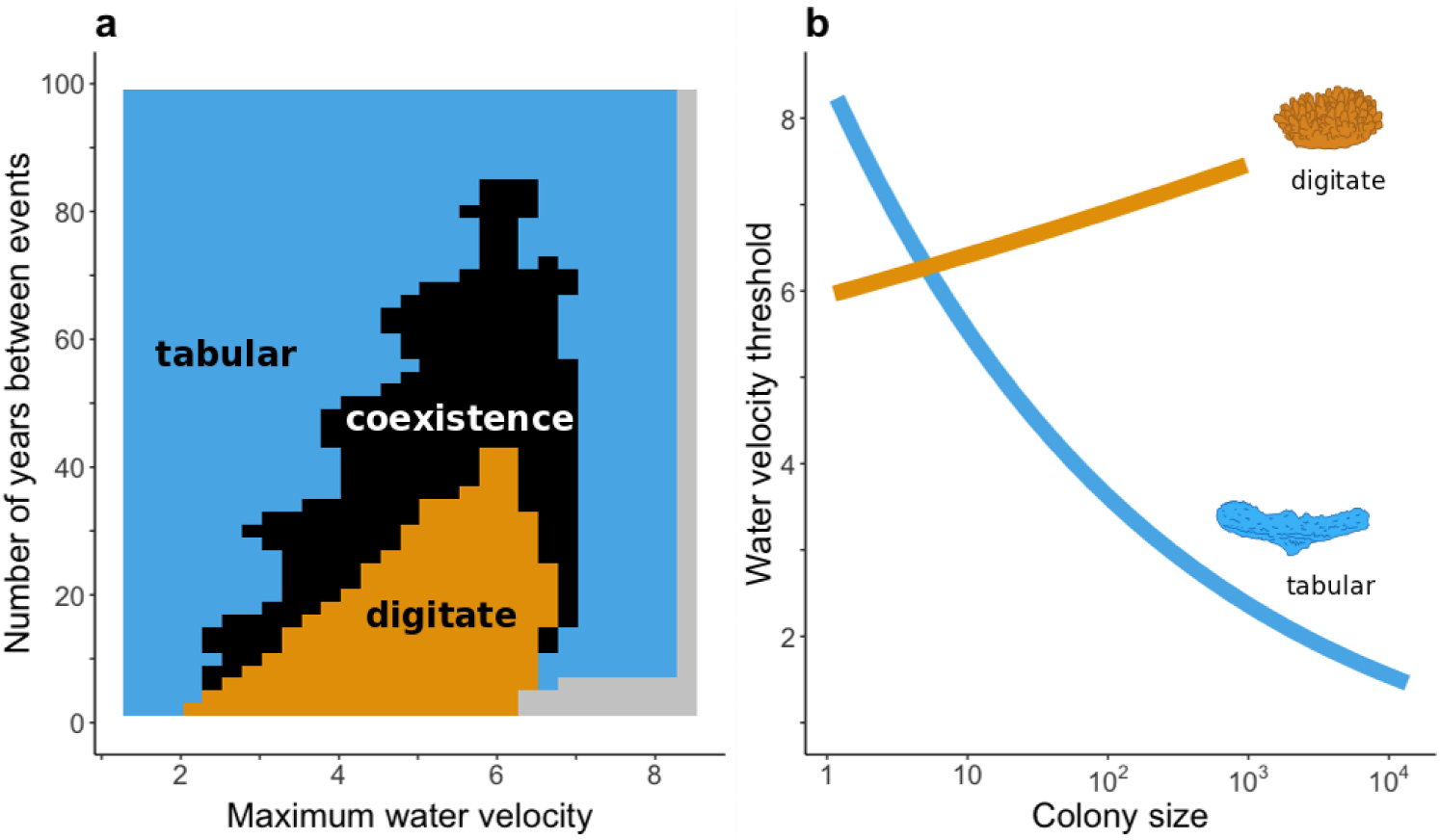
Effect of frequency and intensity of disturbance on species coexistence. Panel a- Competitive outcomes depending on the intensity (maximum water velocity; ms^-1^) and frequency (number of years between disturbance events) of disturbance. Colours indicate combinations of competitive outcomes: the digitate population outcompetes the tabular population in orange areas, the tabular population outcompetes the digitate population in blue areas, in grey areas both competitors go extinct, and coexistence is possible in black areas. Panel b- Minimum water velocity required to dislodge a colony depending on colony size (cm^2^) (estimates are from (Madin *et al.* 2014)). The orange line shows the relationship for the digitate colonies and the blue line shows the relationship for the tabular colonies.

The contribution of nonlinear responses of population growth rates to coexistence has received considerably less attention than the contribution of nonadditive responses (mainly the storage effect). Nonlinear responses to coexistence were initially thought to be more limited than for the latter (Chesson 1994), although recent studies suggest otherwise (Letten *et al.* 2018; Hallett *et al.* 2019; Zepeda & Martorell 2019). Importantly, the contribution of nonlinear responses to coexistence was thought to be limited to competition (e.g., (Chesson 1994, 2000)) until very recently (Ellner *et al.* 2019). In our model, fluctuations in competition were limiting, rather than promoting, coexistence (Fig. 2-a). Since coral demographic rates are tightly linked with colony morphology (Madin *et al.* 2014; Álvarez-Noriega *et al.* 2016), we expect our results to hold for competition between any species of these morphologies. Tabular and digitate corals of the genus *Acropora* have a widespread distribution across the Indo-Pacific and are abundant in wave-exposed reef environments (Done 1982). More broadly, our findings highlight the potential for nonlinear responses in population growth rate to promote coexistence whenever differences in mechanical stability produce differential fluctuations in size structure, as long as those fluctuations disadvantage the population with the higher intrinsic growth rate under average conditions. Because overtopping growth forms tend to be both fast-growing and mechanically unstable (Jackson 1979), nonlinearities are likely to play a role in competition involving species with these growth forms.

Disturbance has long been thought to be an important contributor to coral species coexistence (Connell 1978), but the lack of a mechanism has made this idea controversial in recent years (Chesson & Huntly 1997; Fox 2013). We show that environmental fluctuations can promote coexistence of species that differ in their size-dependent susceptibility to disturbance, limiting the ability of superior competitors to form monocultures and exclude inferior competitors. While other mechanisms are likely to operate to promote coexistence in coral assemblages (including spatial heterogeneity (Hoogenboom *et al.* 2011) and asymmetries and fluctuations in metapopulation connectivity (Salomon *et al.* 2010)), the fact that hydrodynamic disturbances’ transient effects on coral assemblages are large in magnitude and consistent with the model analysed here suggests that the contribution made by hydrodynamic disturbance to coexistence in wave-exposed habitats with fast-growing, top-heavy coral species might be substantial. If so, anthropogenic changes that alter hydrodynamic disturbance regimes (Knutson & Tuleya 2004) or species’ skeletal densities and therefore their vulnerability to mechanical disturbance (Madin *et al.* 2012) are likely to affect coral assemblages in ways that will not be captured by commonly used methods of projecting reef futures, which either consider coral cover to be a single population, or which aggregate the projections of single-species models. Moreover, size-dependent responses to the environment are common in nature (Tredennick *et al.* 2018), and are likely to vary among species. For instance, trees differ in their size-dependent response to drought (Zang *et al.* 2012) and size-dependent fishing pressure affects fish species differently (Genner *et al.* 2010). Consequently, effects of episodic mortality agents on coexistence via nonlinearities in population growth rates are likely to be more widespread in nature than currently recognised.

## Methods

### Analysis overview

First, we specified the competition model and parameterised demographic rates and the disturbance regime. Then, we did an invasibility analysis in the presence of environmental fluctuations to test whether coexistence was possible, and we compared it to an invasibility analysis in a constant environment to investigate if coexistence was mediated by environmental fluctuations. After determining that coexistence was fluctuation-dependent, we decomposed population growth rate into the contribution of: 1) demographic rates at constant (average) conditions, 2) fluctuating size structures, 3) fluctuating competition (proportion of occupied space), and 4) joint fluctuation of competition and size structures. Finally, to test whether competitive outcomes were dependent on the strength and frequency of the disturbance regime, we did invasibility analyses over a range of wind intensities and frequencies. See Figure S1 for a diagrammatic summary of our approach. We explain each step in detail below.

### Competitive model specification

We used integral projection models (Easterling *et al.* 2000) (IPMs) to characterise community dynamics. In the model, demographic processes over each year were divided into two sub-intervals: from time *t*-1 to time *t-h* (where *h* ∈ [1 − *z*, 1) and *z* → 0), when reproduction and larval settlement occurred, and 2) from time *t-h* to time *t*, when disturbance, growth, and survival occurred. In other words, reproduction occurred before growth and survival. We adopted this approach because settlement usually peaks shortly (1-2 weeks) after spawning (Miller & Mundy 2003; Nozawa & Harrison 2008), and thus most growth and mortality of established corals would occur outside this interval. Because coral recruitment is proportional to unoccupied space(Connell *et al.* 1997), we modelled the proportion of larvae successfully recruiting as depending linearly on free space availability 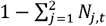, where *N*_*j,t*_ is the proportion of space occupied by species *j* at time *t. N*_*j,t*_ was calculated by integrating the density of colonies of size *y* at time *t* (*n*_*j*_(*y, t*)) times their planar area, and then normalizing by the total habitat area (*A*):

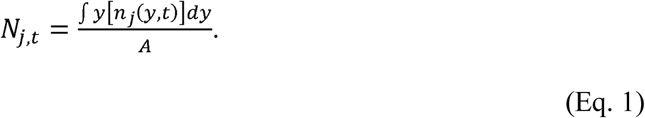

The density of colonies of size *x* at time *t-h* for species *j* (*n*_*j*_(*x, t* − *h*)) was the sum of (i) the density of colonies of size *x* just before settlement (*n*_*j*_(*x, t* − 1)) and (ii) the density of successful settlers of size *x* at time *t-h* produced through reproduction of colonies of size *x* at time *t*-1. The number of possible settlers was given by the integral of the fecundity kernel (*F*_*j*_(*x, x* ′)) times the size distribution at time *t*-1 (*n*_*j*_(*x, t* − 1)). The fecundity kernel was a surface containing transitions from a parent of size *x* at time *t*-1 to an offspring of size *x’* at time *t-h*, and thus it implicitly included reproductive output, larval survival, and successful settlement. With the above assumptions, the density of successful settlers was:

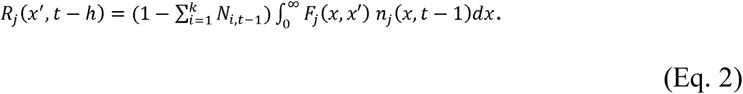

The size distribution at time *t-h* was then:

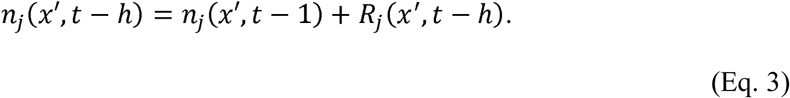

In the second sub interval (*t-h* to time *t*), the model predicted growth and survival of all corals (including those newly recruited) and the proportion of space occupied by each species was calculated (*N*_*j*_). Survival from time *t-h* to time *t* (*S*_*j,t*−*h*_(*x*′)) had two components: a stochastic component that depends on susceptibility to the strongest yearly mechanical disturbance (*D*_*j,t* −*h*_(*x*′)), and a deterministic component that represents ‘background’ mortality (*M*_*j*_ (*x*′); i.e. mortality independent of mechanical disturbance):

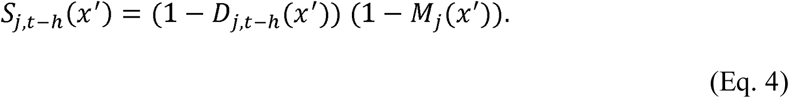

The distribution of colonies of size *y* at time *t*+*1* depended on the survival of colonies of size *x* (*S*_*j, t*−*h*_ (*x*)) and their size-dependent growth to colony size *y* (*G*_*j*_ (*x, y*)) from time *t-h* to time *t:*

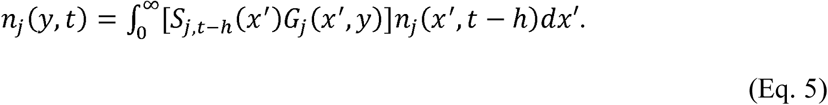

We modelled growth as density-independent, because no effect of competition on colony growth could be detected for the modelled species at the site (Álvarez-Noriega *et al.* 2018). Specifically, growth was modelled as a linear function of size on a logarithmic scale (*y*_*i*_∼*β*_0_ + *β*_1_*x*′_*i*_ + *ε*_*i*_) using published data from the study site (Dornelas *et al.* 2017).

### Parameter estimation

Model parameters are defined in the extended data (Table S1).

Growth, fecundity, and background mortality were obtained from a 5-yr data set of 30 colonies per species on the reef crest of Lizard Island, northern Great Barrier Reef (14.699839°S, 145.448674°E). Parameter estimates were obtained from previously-published analyses of these data (Madin *et al.* 2014; Álvarez-Noriega *et al.* 2016; Dornelas *et al.* 2017), and are reported in the **R** scripts for this paper, which will be publicly available in Github upon publication. For each growth form, demographic data of two species -*Acropora hyacinthus* and *Acropora cytherea* for tabular corals and *Acropora cf. digitifera* and *Acropora humilis* for digitate corals-were pooled for analysis (Fig. S2). *Acropora hyacinthus* and *A. cf. digitifera* were among the most abundant coral species at the site (Dornelas & Connolly 2008). Previous analyses indicate that growth, mortality, and fecundity all vary substantially more between growth forms than between species of the same growth form (Madin *et al.* 2014; Álvarez-Noriega *et al.* 2016; Dornelas *et al.* 2017).

The fecundity function (*F*_*j*_ (*x, x*′)) for species *j* depended on: the size-dependent probability of a polyp being mature (*p*_*j,x*_), the size-dependent number of oocytes per mature polyp (*m*_*j,x*_), the number of polyps per projected unit area (*ρ*_*j*_), the projected area of the colony (*a*_*j*_) and the settlement probability (*q*_*j*_):

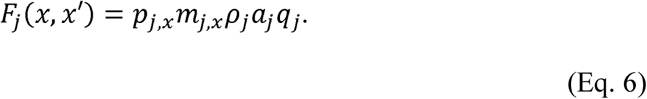

The settlement probability included the probability of an egg being fertilised and becoming a larva, and the probability of that larva successfully settling. Since larval mortality is density-independent and per-capita settlement, if affected at all, is positively affected by higher larval densities(Heyward *et al.* 2002; Edwards *et al.* 2015; Doropoulos *et al.* 2017, 2018), we assumed no competition among larvae (i.e., successful recruitment depended on unoccupied space, but not on the density of offspring seeking to settle).

All terms in eq. (6) except for settlement probability were obtained from Álvarez-Noriega et al. (2016)(Álvarez-Noriega *et al.* 2016). Since there is no information available on settlement probability, we fixed a value resulting in average coral cover of about 75% of each morphology in the absence of competitors and in the presence of disturbance. However, we considered a range of values of this parameter to ensure that settlement probabilities resulting in lower average coral cover yielded consistent results (i.e., coexistence by RNC in the presence of environmental fluctuations; no coexistence possible without fluctuations).

Mortality due to mechanical disturbance was determined by comparing each colony’s ‘Colony Shape Factor’ (CSF) to the ‘Dislodgment Mechanical Threshold’ (DMT) imposed by the yearly maximum hydrodynamic disturbance. Both quantities were derived by Madin and Connolly (Madin & Connolly 2006b), and field-tested on Lizard Island, at a reef adjacent to our study site. When the DMT of an event exceeds the CSF of a colony, the colony is predicted to be dislodged.

In each year of the simulation, a random wind velocity was drawn from a gamma distribution (α=2.18, β=0.35), with parameters estimated from the distribution of a 37-year wind velocity data for the Low Isles (16.383°S, 145.567°E) (from the Australian Bureau of Meteorology), approximately 180 km south of Lizard Island (Madin *et al.* 2006). Water velocity at the reef crest as a function of wind velocity was estimated using wind and water velocity at the reef crest collected at the site(Madin *et al.* 2006). We predicted water velocity, *u*, as a saturating function of wind velocity, *v*, because wave energy is limited by fetch and depth:

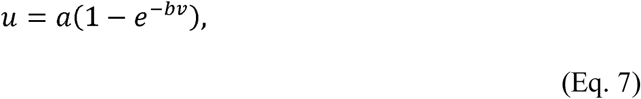

where *a* and *b* are fitted parameters, estimated by least-squares estimation (*a=* 5.10, *b=* 0.04; Fig. S3). Using previously calibrated relationships between colony size and CSF for our study species at Lizard Island (Madin *et al.* 2014), colonies were predicted to dislodge if the DMT imposed by the yearly maximum wind velocity was smaller than the colony’s estimated CSF (Madin & Connolly 2006b). Since dislodged colonies have very low survival rates (Smith & Hughes 1999), colony dislodgement was assumed to cause colony mortality.

Background mortality (mortality independent of mechanical disturbance) was estimated from mortality data from 2009-2012 (Madin *et al.* 2014). CSF for each individual colony was estimated and compared to the maximum dislodgement mechanical threshold imposed by the environment that year (estimated from wind data at the site(Australian Institute of Marine Science 2017)). As predicted by theory (Madin & Connolly 2006b) and validated with mortality data on the reef (Madin *et al.* 2014), colonies that had CSF larger than the DMT estimated for that year were assumed to have been dislodged. Colonies predicted to dislodge were assumed dead and were removed from the data set from which background mortality was estimated. With the remaining data, two linear models with a binomial error structure were fitted for each growth form: one with colony area as an explanatory variable (log-scale) and one independent of colony area. Models were compared using AIC, and the best-fit model for each growth form was used in the simulations (Tables S2-S3; Figs. S4-S5).

### Analysis of coexistence

Partitioning the contribution of the fluctuation-dependent mechanisms using analytical solutions for quadratic approximations (Chesson 1994) is unfeasible in complex models (e.g. multiple-step, stage/size-dependent models); therefore, we estimated terms needed for the analytical solution via simulations.

Overall, the finite rate of increase in cover of species *j* between time *t-1* and time *t* (*λ*_*j*_ (*t*)) in terms of proportion of occupied space (i.e. the factor by which the proportion of space occupied by species *j* changes) is:

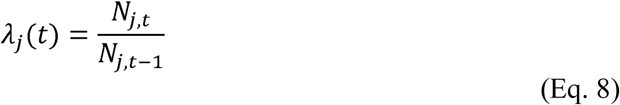

and the overall specific population growth rate (*r*_*j*_ (*t*)) is:

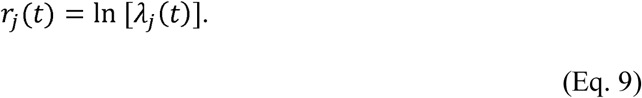

### Decomposing growth rate into contributions from different sources of variation

We follow Ellner et al. (Ellner *et al.* 2019) to decompose mean population growth rate of each species 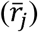 into population growth rate at mean conditions 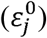, the contribution of fluctuations in competition 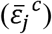, fluctuations in size-structure 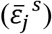, and the interaction between fluctuating competition and size-structure 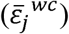:

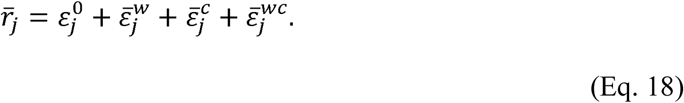

The approach involves computing population growth under different scenarios: one where all varying factors are allowed to fluctuate, and one for each possible combination of one of more of the varying factors fixed at its mean value from the stochastic simulations. The population growth rates from the different scenarios are compared to quantify the contribution of each factor varying on its own and in combination with other factors.

First, a simulation was run in which one population was introduced at a coral cover <10^−9^ in a system where the competitor had been resident for 2000 years; the simulation ran for 300 years after the invasion. In this first simulation (the baseline simulation), wind velocities were randomly drawn from the gamma distribution fitted to the wind velocity data sat Low Isles; from this simulation we estimated 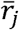 as the mean population growth of species *j* from 50 years after the invasion to the end of the simulations. Competition (*c*), and size-structure (*w*) (both *n* (*x*′, *t* − *h*) and *n*(*y,t* + 1)) were recorded for each year of the simulation (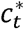 and 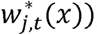, where *w*_*j,t*_ (*x*) is the size distribution of colony sizes of species *j* at time *t* (normalized to integrate to unity) and *c*_*t*_ is the proportion of free space at time t; the asterisks indicates values for the baseline simulation. Using these values, we then calculated their temporal averages 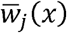 and 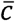 and ran a simulation in which *w*_*j,t*_(*x*) and *c*_*t*_ were fixed at those mean values to estimate 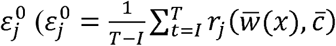, where *I* is time after 50 years after the invader was introduced (*I=*2050; to be consistent with the previous growth rate decomposition) and *T* is time at the end of the simulations (*T*=2300); i.e. the mean population growth calculated for this set of simulations). To maintain a constant size structure, the total number of colonies of species *j* at time *t* were summed and the corresponding proportion was allocated to each size to match the temporal average size structure from the stochastic simulations 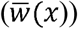. Here, the proportion of free space was fixed at 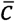 during all years of the simulation. To calculate 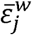, another simulation was run where 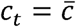 but the size-structures were set at the value recorded for the baseline simulation 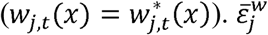 was the effect of fluctuations in size-structure on population growth, independent of the effect of mean size-structure 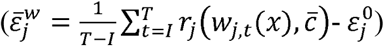. Similarly, 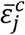 (where 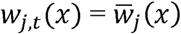 and 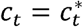) was estimated as the effect of competition independent of the effect of its mean (Table S4 for all the relevant formulas). The interaction between *w* and 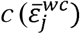, was the effect of both *w*_*j,t*_ (*x*) and *c*_*t*_ varying together 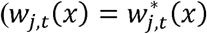 and 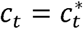, independent of the effects of mean size-structure and mean competition, and of size-structure and competition fluctuating on their own: 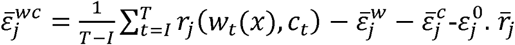 was then defined according to Eq. (18).

### Sensitivity analysis: Coexistence under different disturbance frequency-intensity regimes

To investigate the combinations of wave disturbance frequency and intensity that allowed tabular and digitate populations to coexist, we simulated invasions of both competitors under a range of water velocities (i.e. disturbance intensity) with a range of years between disturbances (frequencies). Water velocity ranged from 1.4 (no colonies are dislodged) to 8.5 ms^-1^ (all colonies are dislodged), in 0.25ms^-1^ increments; the number of years between disturbances ranged from zero to 100 years in increments of 2 years. Before each invasion, 2000 years of simulations were run to allow the resident to reach a stable range of coral cover; after each invasion, 300 years more of community dynamics were simulated. Coexistence was possible if both competitors -as invaders- had a higher mean proportion of space cover in the last 100 years of the simulation than in the 100 years following the invasion 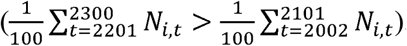. If this was true for the tabular population, but not for the digitate, the tabular population was assumed to dominate; conversely, the digitate population was assumed to dominate when it was able to invade a tabular population, but the tabular population was not able to invade a digitate resident. See supplementary material for a stochastic version of this analysis.

All simulations were done in R (R Core Team 2018) (version 3.5.2).

## Acknowledgments

We thank R. Aronson, P. Adler, and A. Álvarez-Noriega for helpful comments on an earlier version of this work, and D. Falster for valuable advice.

## Author contributions

All authors conceived the study. JSM provided the data to estimate the wind-water velocity relationship at the site and helped estimate ‘background mortality’. MAN did the simulations and wrote the first draft with help from SRC. All authors contributed to subsequent drafts.

## Data and materials availability

All scripts will be publicly available in Github upon publication. No new data were used.

## References

Adler, P.B., HilleRisLambers, J., Kyriakidis, P.C., Guan, Q. & Levine, J.M. (2006). Climate variability has a stabilizing effect on the coexistence of prairie grasses. Proc. Natl. Acad. Sci. U. S. A., 103, 12793–8.

Álvarez-Noriega, M., Baird, A.H., Dornelas, M., Madin, J.S. & Connolly, S.R. (2018). Negligible effect of competition on coral colony growth. Ecol..

Álvarez-Noriega, M., Baird, A.H., Dornelas, M., Madin, J.S., Cumbo, V.R. & Connolly, S.R. (2016). Fecundity and the demographic strategies of coral morphologies. Ecology, 97, 3485–3493.

Angert, A.L., Huxman, T.E., Chesson, P. & Venable, D.L. (2009). Functional tradeoffs determine species coexistence via the storage effect. Proc. Natl. Acad. Sci. U. S. A., 106, 11641–5.

Armstrong, R.A. & McGehee, R. (1980). Competitive exclusion. Am. Nat.

Aronson, R. & Precht, W. (1995). Landscape patterns of reef coral diversity: A test of the intermediate disturbance hypothesis. J. Exp. Mar. Bio. Ecol., 192, 1–14.

Australian Institute of Marine Science. (2017). Lizard Island wind speed. Https://data.gov.au/dataset/imos-faimms-sensor-network-data-lizard-island-weather-station-wind-speed-from-13-aug-2010-20171,.

Baird, A. & Hughes, T. (2000). Competitive dominance by tabular corals: an experimental analysis of recruitment and survival of understorey assemblages. J. Exp. Mar. Bio. Ecol., 251, 117–132.

Bellwood, D.R. & Hughes, T.P. (2001). Regional-scale assembly rules and biodiversity of coral reefs. Science (80-.).

Cáceres, C.E. (1997). Temporal variation, dormancy, and coexistence: a field test of the storage effect. Proc. Natl. Acad. Sci. U. S. A., 94, 9171–9175.

Chesson, P. (1994). Multispecies competition in variable environments. Theor. Popul. Biol.

Chesson, P. (2000). Mechanisms of maintenance of species diversity. Annu. Rev. Ecol. Syst..

Chesson, P. & Huntly, N. (1997). The roles of harsh and fluctuating conditions in the dynamics of ecological communities. Am. Nat.

Chesson, P.L. & Warner, R.R. (1981). Environmental variability promotes coexistence in lottery competitive systems. Am. Nat.

Connell, J.H. (1978). Diversity in tropical rain forests and coral reefs. Science (80-.).

Connell, J.H., Hughes, T.P. & Wallace, C.C. (1997). A 30-year study of coral abundance, recruitment, and disturbance at several scales in space and time. Ecol. Monogr., 67, 461–488.

Connell, J.H., Hughes, T.P., Wallace, C.C., Tanner, J.E., Harms, K.E. & Kerr, A.M. (2004). A long-term study of competition and diversity of corals. Ecol. Monogr.

Done, T.J. (1982). Patterns in the distribution of coral communities across the central Great Barrier Reef. Coral Reefs.

Dornelas, M. & Connolly, S.R. (2008). Multiple modes in a coral species abundance distribution. Ecol. Lett., 11, 1008–16.

Dornelas, M., Madin, J.S., Baird, A.H. & Connolly, S.R. (2017). Allometric growth in reef-building corals. Proceedings. Biol. Sci.

Doropoulos, C., Evensen, N.R., Gómez-Lemos, L.A. & Babcock, R.C. (2017). Density-dependent coral recruitment displays divergent responses during distinct early life-history stages. R. Soc. open Sci., 4, 170082.

Doropoulos, C., Gómez-Lemos, L. & Babcock, R. (2018). Exploring variable patterns of density-dependent larval settlement among corals with distinct and shared functional traits. J. Int. Soc. Reef Stud., 37, 25–29.

Easterling, M.R., Ellner, S.P. & Dixon, P.M. (2000). Size-specific sensitivity: Applying a new structured population model. Ecology, 81.

Edwards, A.J., Guest, J.R., Heyward, A.J., Villanueva, R.D., Baria, M.V., Bollozos, I.S.F., et al. (2015). Direct seeding of mass-cultured coral larvae is not an effective option for reef rehabilitation. Mar. Ecol. Prog. Ser., 525, 105–116.

Ellner, S.P., Snyder, R.E., Adler, P.B. & Hooker, G. (2019). An expanded modern coexistence theory for empirical applications. Ecol. Lett., 22, 3.

Fox, J.W. (2013). The intermediate disturbance hypothesis should be abandoned. Trends Ecol. Evol.

Genner, M.J., Sims, D.W., Southward, A.J., Budd, G.C., Masterson, P., McHugh, M., et al. (2010). Body size-dependent responses of a marine fish assemblage to climate change and fishing over a century-long scale. Glob. Chang. Biol.

Hallett, L.M., Shoemaker, L.G., White, C.T. & Suding, K.N. (2019). Rainfall variability maintains grass-forb species coexistence. Ecol. Lett., 22, 1658–1667.

Heyward, A.J., Smith, L.D., Rees, M. & Field, S.N. (2002). Enhancement of coral recruitment by in situ mass culture of coral larvae. Mar. Ecol. Prog. Ser., 230, 113–118.

Hoogenboom, M., Connolly, S. & Anthony, K. (2011). Biotic and abiotic correlates of tissue quality for common scleractinian corals. Mar. Ecol. Prog. Ser., 438, 119–128.

Hutchinson, G.E. (1961). The paradox of the plankton. Am. Nat.

Jackson, J.B.C. (1979). Morphological strategies of sessile animals. Biol. Syst. Colon. Org. Academic Press, London.

Jensen, J.L.W. V. (1906). Sur les fonctions convexes et les inégalités entre les valeurs moyennes. Acta Math..

Knutson, T.R. & Tuleya, R.E. (2004). Impact of CO_2_-induced warming on simulated hurricane intensity and precipitation sensitivity to the choice of climate model and convective parameterization. J. Clim., 17, 3477–3495.

Letten, A.D., Dhami, M.K., Ke, P.J. & Fukami, T. (2018). Species coexistence through simultaneous fluctuation-dependent mechanisms. Proc. Natl. Acad. Sci. U. S. A.

Madin, J.S., Baird, A.H., Dornelas, M. & Connolly, S.R. (2014). Mechanical vulnerability explains size-dependent mortality of reef corals. Ecol. Lett., 17, 1008–1015.

Madin, J.S., Black, K.P. & Connolly, S.R. (2006). Scaling water motion on coral reefs: from regional to organismal scales. Coral Reefs, 25, 635–644.

Madin, J.S. & Connolly, S.R. (2006a). Ecological consequences of major hydrodynamic disturbances on coral reefs. Nature, 444, 477–80.

Madin, J.S. & Connolly, S.R. (2006b). Ecological consequences of major hydrodynamic disturbances on coral reefs. Nature, 444, 477–80.

Madin, J.S., Hughes, T.P. & Connolly, S.R. (2012). Calcification, storm damage and population resilience of tabular corals under climate change. PLoS One, 7, e46637.

Massel, S.R. & Done, T.J. (1993). Effects of cyclone waves on massive coral assemblages on the Great Barrier Reef: meteorology, hydrodynamics and demography. Coral Reefs, 12, 153–166.

Miller, A.D., Roxburgh, S.H. & Shea, K. (2011). How frequency and intensity shape diversity-disturbance relationships. Proc. Natl. Acad. Sci. U. S. A., 108, 5643–8.

Miller, K. & Mundy, C. (2003). Rapid settlement in broadcast spawning corals: implications for larval dispersal. Coral reefs, 22, 99–106.

Muko, S., Arakaki, S., Nagao, M. & Sakai, K. (2013). Growth form-dependent response to physical disturbance and thermal stress in Acropora corals. Coral Reefs.

Nozawa, Y. & Harrison, P.L. (2008). Temporal patterns of larval settlement and survivorship of two broadcast-spawning acroporid corals. Mar. Biol., 155, 347–351.

R Core Team. (2018). R: A Language and Environment for Statistical Computing. R Foundation for Statistical Computing.

Roxburgh, S.H., Shea, K. & Wilson, J.B. (2004). The intermediate disturbance hypothesis: patch dynamics and mechanisms of species coexistence. Ecology.

Salomon, Y., Connolly, S.R. & Bode, L. (2010). Effects of asymmetric dispersal on the coexistence of competing species. Ecol. Lett.

Smith, L.D. & Hughes, T.P. (1999). An experimental assessment of survival, re-attachment and fecundity of coral fragments.

Tanner, J.E., Hughes, T.P. & Connell, J.H. (1994). Species coexistence, keystone species, and succession: a sensitivity analysis. Ecology.

Tredennick, A.T., Teller, B. j., Adler, P.B., Hooker, G. & Ellner, S.P. (2018). Size-by-environment interactions: a neglected dimension of species’ responses to environmental variation. Ecol. Lett., 21, 1757–1770.

Zang, C., Pretzsch, H. & Rothe, A. (2012). Size-dependent responses to summer drought in Scots pine, Norway spruce and common oak. Trees.

Zepeda, V. & Martorell, C. (2019). Fluctuation-independent niche differentiation and relative non-linearity drive coexistence in a species-rich grassland. Ecol..

