## Supplementary Information for "Disturbance-induced changes in size-structure promote coral biodiversity"

This PDF file includes:

Supplementary text

Figures S1 to S7

Tables S1 to S4

SI References

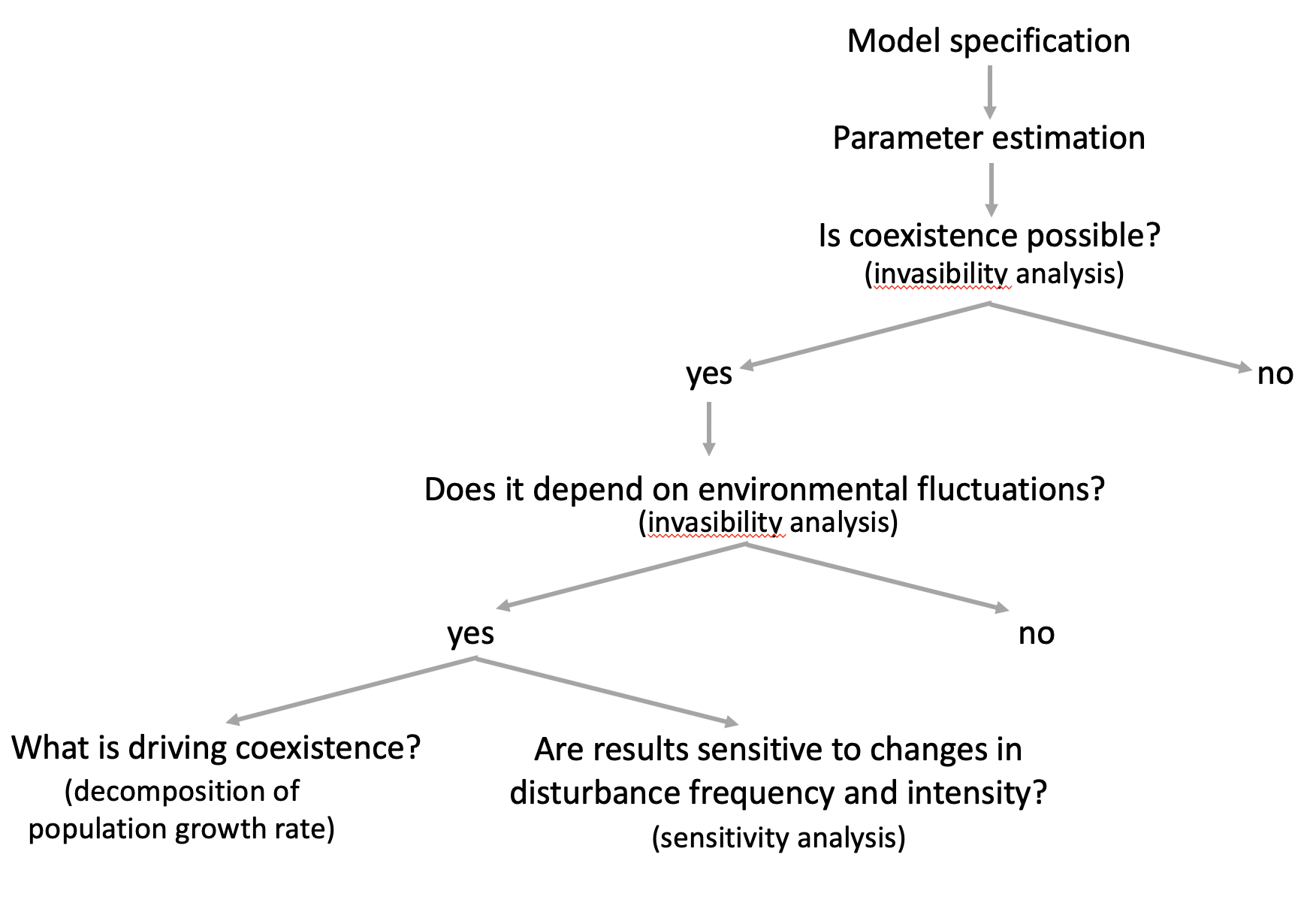

**Figure S1.** Diagram summarising the steps of the analysis.

**Table S1.** Model parameters.

| $N_{j,t}$ | proportion of space occupied by species *j* at time *t* |
| --- | --- |
| $n_{j}\left( y,t \right)$ | density of colonies of species *j* size *y* at time *t* |
| *A* | total habitat area |
| $F_{j}\left( x,x' \right)$) | fecundity function of species *j* |
| $R_{j}\left( x^{'},t-h \right)$ | density of successful settlers of species *j* |
| $S_{j,t-h}\left( x^{'} \right)$ | survival of colonies size *x’* of species *j* from time *t-h* to time *t* |
| $D_{j,t-h}\left( x^{'} \right)$ | mortality of colonies size *x’* of species *j* from time *t-h* to time *t* caused by mechanical disturbance |
| $M_{j}\left( x^{'} \right)$ | mortality of colonies size *x’* of species *j* from time *t-h* to time *t* independent of mechanical disturbance |
| $G_{j}\left( x, y \right)$ | growth function of species *j* |
| $p_{x}$ | probability of a polyp being mature from a colony size *x* |
| $m_{x}$ | number of oocytes per mature polyp of a colony size *x* |
| $\rho$ | number of polyps per projected unit area |
| *w* | projected area of the colony |
| *q* | settlement probability |

**Table S2.** Comparisons of binomial linear models predicting disturbance-independent survival of tabular and digitate colonies with and without colony area (log-scale, cm^2^) as explanatory variables.

| Competitor | Model | AIC | AIC weights |
| --- | --- | --- | --- |
| tabular | log-area | 41.72 | 0.32 |
|  | - | 40.19 | 0.68 |
| digitate | log-area | 136.61 | 0.81 |
|  | - | 139.51 | 0.19 |

**Table S3.** Intercept and slope estimates for the logit transform of disturbance-independent survival probability as a function of colony area (log-scale, cm^2^).

| Competitor | Effect | Estimate | Std. Error | z value | Pr(>\|z\|) |
| --- | --- | --- | --- | --- | --- |
| tabular | (intercept) | 0.693 | 0.387 | 1.790 | 0.074 |
|  | log-area | - | - | - | - |
| digitate | (intercept) | -0.409 | 1.231 | -0.332 | 0.740 |
|  | log-area | 0.525 | 0.235 | 2.234 | 0.026 |

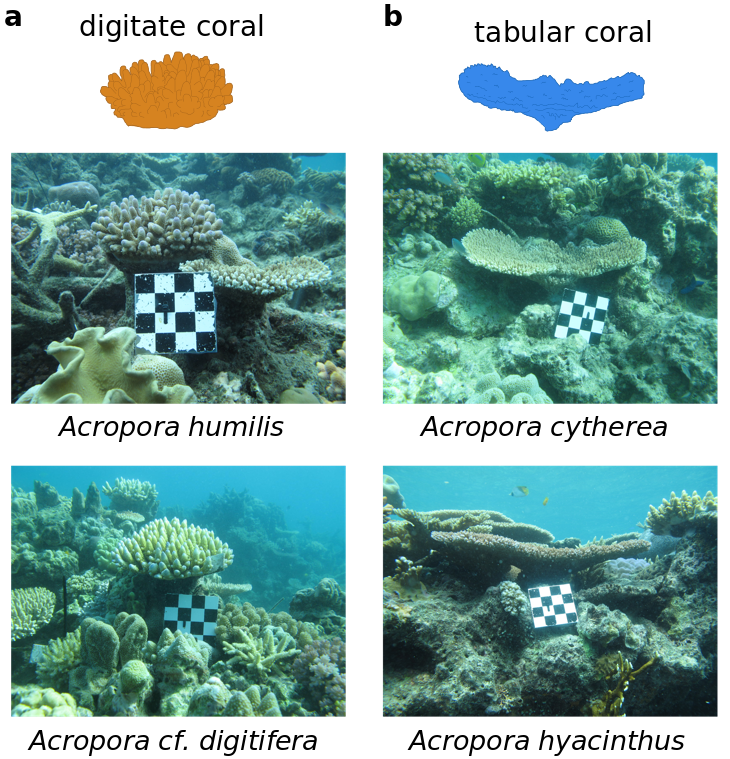

**Figure S2.** Panel a- Stylized illustrations of a digitate colony. Panel b- Stylized illustrations of a tabular colony. The species representing each morphology are listed below each illustration with the respective side-on photo of the species at the site.

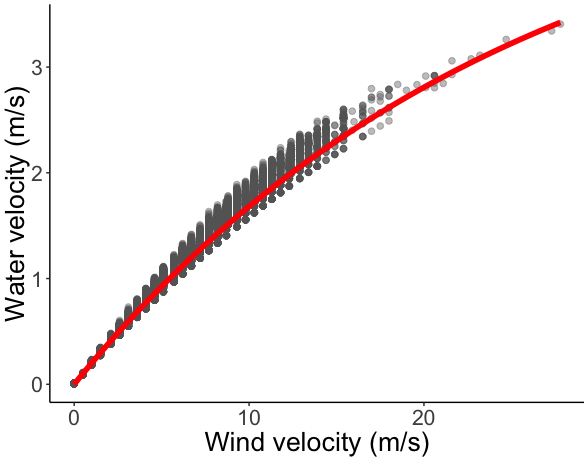

**Figure S3**. Fit model predicting water velocity ($u$; ms^-1^) as a saturated function of wind velocity ($v$; ms^-1^) ($u=5.10\left( 1-e^{-0.04v} \right))$. The red line shows the fitted model and the grey points show the data.

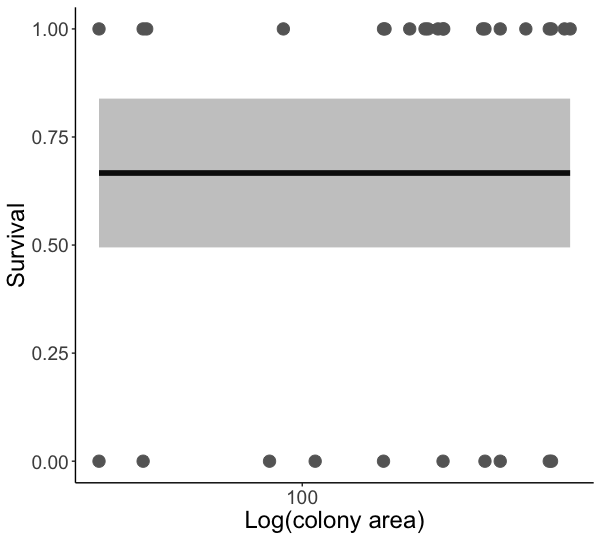

**Figure S4.** Predictions of the best-fit model predicting disturbance-independent survival of tabular colonies vs. colony size (log-scale, cm^2^). The solid line shows model predictions, the grey ribbon shows standard errors, and the solid circles show the data points.

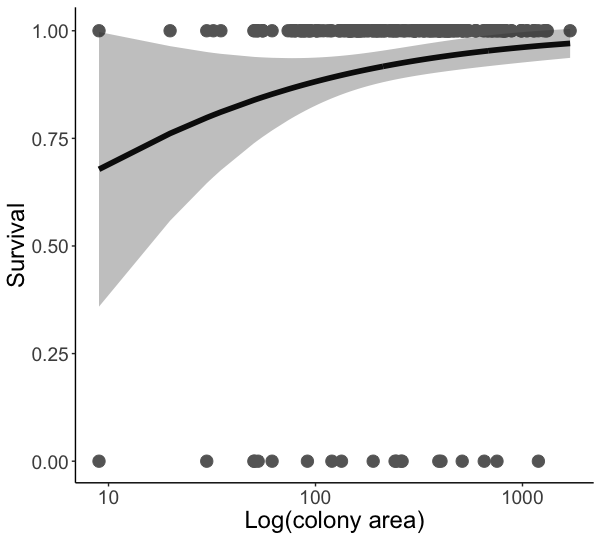

**Figure S5**. Predictions of the best-fit model predicting disturbance-independent survival of digitate colonies vs. colony size (log-scale, cm^2^). The solid line shows model predictions, the grey ribbon shows standard errors, and the solid circles show the data points.

**Table S4.** Formulas for the decomposition of growth rate into contributions from different sources of variation following Ellner et al. (2019)(Ellner *et al.* 2019). $w_{x,t}$ and $c_{t}$, take their values in the baseline simulation ($w_{x,t}^{*}$ and $c_{t}^{*}$).

| Term in Fig. 2 | Term in Ellner et al. (2019) | Formula |
| --- | --- | --- |
| *constant* | $\varepsilon_{0}$ | $r_{j}(\bar{w}(x)$, $\bar{c}$) |
| *size structure* | $\bar{\varepsilon}^{w}$ | $\frac{1}{T-I}\sum_{t=I}^{T} r_{j}\left( {w(x)}_{t}, \bar{c}) \right)$-$\varepsilon_{j}^{0}$ |
| *competition* | $\bar{\varepsilon}^{c}$ | $\frac{1}{T-I}\sum_{t=I}^{T} r_{j}\left( \bar{w}(x), c_{t} \right)$-$\varepsilon_{j}^{0}$ |
| *size* ***x*** *competition* | $\bar{\varepsilon}^{wc}$ | $\frac{1}{T-I}\sum_{t=I}^{T} r_{j}\left( {w(x)}_{t}, c_{t} \right)-\bar{\varepsilon}_{j}^{w}-\bar{\varepsilon}_{j}^{c}$-$\varepsilon_{j}^{0}$ |

Nonlinear effects of competition in competitors’ population growth rates

Following Jensen’s inequality (Jensen 1906), the average of the population growth (*r*) as a function of a limiting factor *R* (such as competition) ($\bar{r(R})$) is different to *r* evaluated at the mean of the limiting factor ($r(\bar{R}$)). When the relationship between *r* and *R* is convex, then $\bar{r(R}) \geq r(\bar{R}$)), otherwise $\bar{r(R}) \leq r(\bar{R}$)), which means that nonlinearities in the dependence of a common limiting factor can either boost or depress competitors’ growth rates, relative to a constant environment at mean environmental conditions. When survival and size structure were at their average values but competition fluctuated, *r* ($r\left( \bar{w}(x), c_{t} \right)$) was a concave function of free space for both competitors and thus $r\left( \bar{w}(x), c_{t} \right)$<$r(\bar{w}(x)$, $\bar{c}$) (Fig. S6). However, *r* was more concave -- decreased more rapidly with competition (low free space)-- for the digitate population and in most years the proportion of free space was at a level where $r_{digitate}<r_{tabular}$. These conditions meant that, under fluctuating competition alone, both competitors had a lower *r* than under average conditions, but the reduction in *r* induced by fluctuations was greater for the digitate population, thereby inhibiting coexistence (Fig. 2).

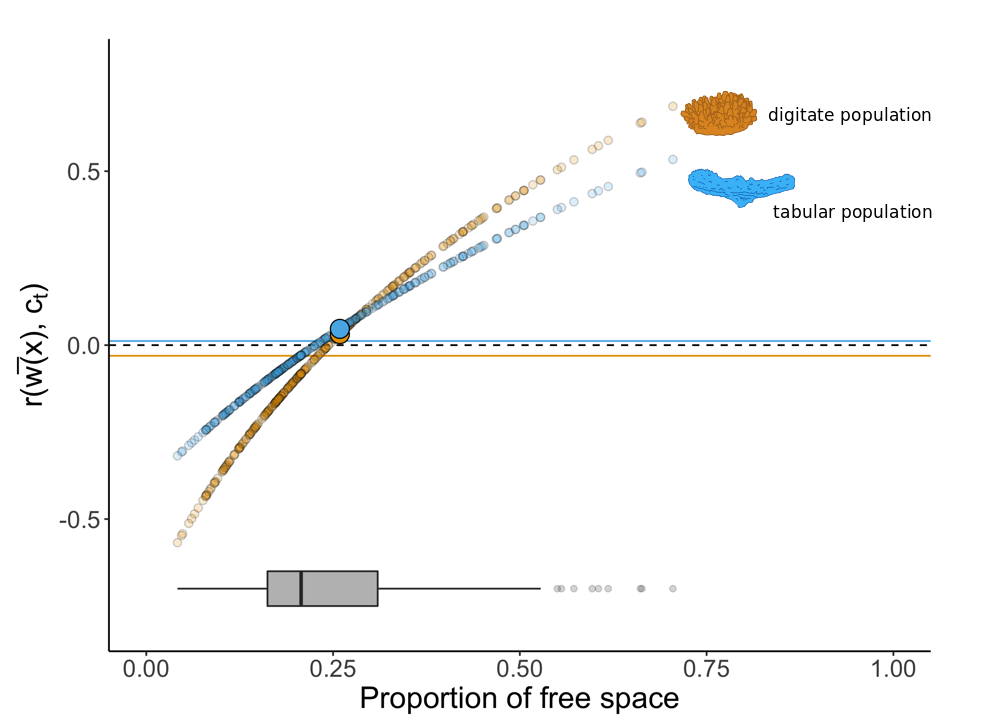

**Figure S6.** Population growth rate when size structures are fixed at their average values but competition fluctuates through time ($r\left( \bar{w}(x), c_{t} \right)$) as a function of the proportion of space available for settlement, in the scenario where the tabular population is at its long-term abundance in the absence of the digitate population, and the digitate population is invading. Blue colours indicate values for the tabular population and orange colours indicate values for the digitate population. The small circles show the relationship between yearly population growth rates and the corresponding proportion of free space (sample for one simulation). The two large circles (one orange and one blue) show the population growth rates for each population at average conditions. The grey horizontal boxplot shows the distribution of values for the proportion of free space. When $r\left( \bar{w}(x), c_{t} \right)=0$ is marked with a black dashed line, the mean values of $r\left( \bar{w}(x), c_{t} \right)$ (i.e., averages in the presence of fluctuations in competition) are marked with a solid horizontal for each population. Note that this average for the digitate population (the orange line) is further below the value under average conditions (the large orange point) than the corresponding quantity for the tabular population.

Sensitivity analysis: Coexistence under different disturbance frequency-intensity regimes (stochastic)

Similar to the sensitivity analysis in the main text, we investigated the combinations of wave disturbance frequency and intensity that allowed tabular and digitate populations to coexist with water velocity ranging from 1.4 (no colonies are dislodged) to 8.5 ms^-1^ (all colonies are dislodged), in 0.25ms^-1^ increments; and the number of years between disturbances ranging from zero to 100 years in increments of 2 years on average. Before each simulation of disturbance intensity *i* and frequency *f,* we assigned a water velocity to each year (either 0 ms^-1^ or *i*), where *i* occurred at a frequency of *f.* In contrast to the deterministic scenario in the main text, we randomly shuffled these values of water velocity so that we would expect disturbance to occur at an average frequency close to *f* but the period between disturbances was not fixed. For each combination of disturbance frequency and intensity, we ran 100 simulations and computed the proportion of those simulations where coexistence was possible. If coexistence did not occur in any simulation, the competitor which was able to invade in most simulations was assigned as the winner. If neither competitor was able to recover from low densities, both competitors were assumed to go extinct. See Figure F7 for the results of this analysis.

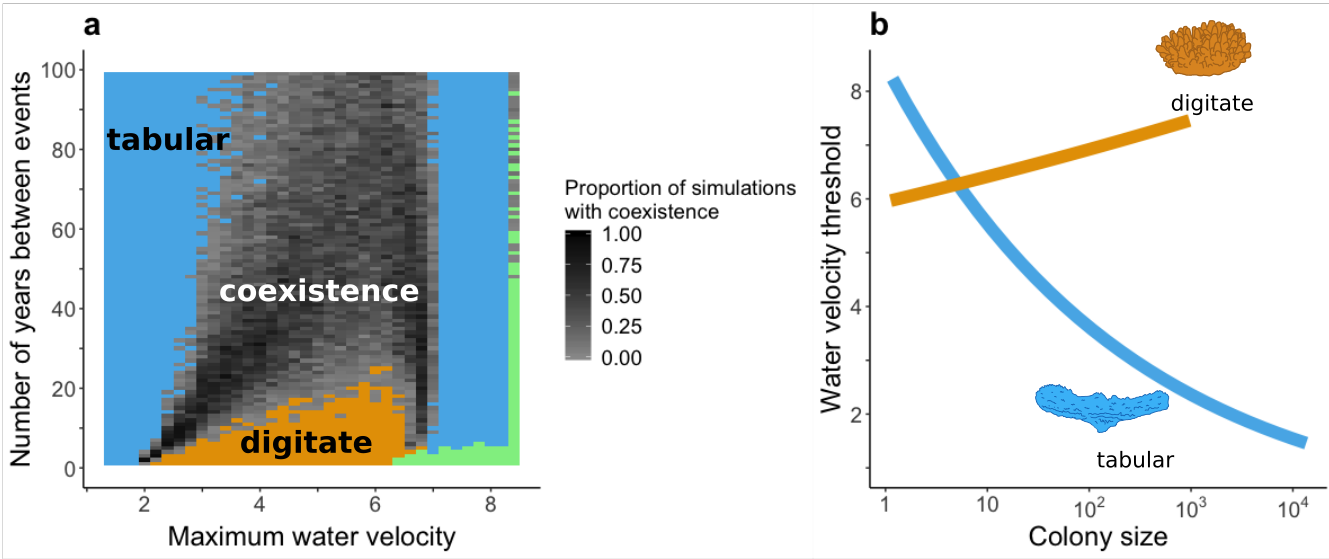

**Figure S7.** Effect of frequency and intensity of disturbance on species coexistence. Panel a- Competitive outcomes depending on the intensity (maximum water velocity; ms^-1^) and expected frequency (number of years between disturbance events) of disturbance for the stochastic scenario. Colours indicate combinations of competitive outcomes: the digitate population outcompetes the tabular population in orange areas, the tabular population outcompetes the digitate population in blue areas, in green areas both competitors go extinct, and coexistence is possible in grey areas. The shade of grey shows the proportion of simulations (out of 100) where coexistence was possible going from light (coexistence only occurred in a small proportion of simulations) to dark grey (coexistence occurred in most simulations) Panel b- Minimum water velocity required to dislodge a colony depending on colony size (cm^2^) (estimates are from (Madin *et al.* 2014)). The orange line shows the relationship for the digitate colonies and the blue line shows the relationship for the tabular colonies.

**References**

Ellner, S.P., Snyder, R.E., Adler, P.B. & Hooker, G. (2019). An expanded modern coexistence theory for empirical applications. *Ecol. Lett.*, 22, 3.

Jensen, J.L.W. V. (1906). Sur les fonctions convexes et les inégalités entre les valeurs moyennes . *Acta Math.* .

Madin, J.S., Baird, A.H., Dornelas, M. & Connolly, S.R. (2014). Mechanical vulnerability explains size-dependent mortality of reef corals. *Ecol. Lett.*, 17, 1008–1015.
